# inGSEA: An Improved Method for Gene Set Enrichment Analysis Using a Weighted Integral Statistic

**DOI:** 10.64898/2026.06.02.729106

**Authors:** Qinyuan Zhang, Qizhai Li

## Abstract

Gene Set Enrichment Analysis (GSEA) is one of the most popular methods for transcriptomic analysis, yet its statistical power is limited when the biological pathways exhibit heterogeneous or non-concordant expression patterns. We propose an improved GSEA method, **in**tegral-based GSEA (inGSEA). inGSEA introduces a novel enrichment score based on the Anderson-Darling weighted integral statistic. The new enrichment score enhances detection power for complex signals, particularly sparse and bidirectional ones, while the Cauchy combination of integral and classic maximum statistics provides robustness across diverse expression patterns. Extensive numerical studies demonstrate that inGSEA achieves superior power and well-calibrated false discoveries. Application to real-world datasets reveals biologically relevant pathways missed by the standard GSEA. inGSEA reduces the computational burden of permutation testing by employing a generalized gamma distribution to approximate the null distribution. inGSEA is accessible as a user-friendly web-based software tool (https://amss-stat.github.io/inGSEA).

## 1 Introduction

In the era of high-throughput transcriptomics, interpreting lists of thousands of differentially expressed genes remains a primary challenge. Rather than focusing on single genes in isolation, Gene Set Analysis (GSA) evaluates predefined groups of genes that share common biological functions or regulatory mechanisms, providing a more interpretable view of transcriptomic states [1, 2]. Among the existing GSA methods, Gene Set Enrichment Analysis (GSEA) [3, 4] has emerged as one of the most popular and influential frameworks, with over 50,000 citations to date. GSEA ranks all genes across the transcriptome based on their correlation with a specific phenotype, and evaluates whether the predefined gene sets are enriched at the extremes of this ranked list. The interpretation of these enrichment results is routinely supported by comprehensive databases, notably the Molecular Signatures Database (MSigDB) [5, 6].

The GSEA framework has inspired numerous methodological extensions and enhancements, such as ssGSEA [7], GSVA [8] and irGSEA [9] for assessing pathway activity at the single-sample level. For the canonical case-control analysis, improvements to GSEA predominantly focus on resolving computational bottlenecks. Tools like fgsea [10] and blitzGSEA [11] substantially improve computational efficiency through optimized permutation algorithms and gamma distribution approximations. Despite these significant computational leaps, the core statistical framework of GSEA has remained unchanged. As noted in the original methodology and GSEA software [4], GSEA assumes that the gene expression changes between two biological states are concordant within the enriched gene set. This behavior is closely tied to the application of a Kolmogorov-Smirnov (KS) type statistic [12] in calculating the Enrichment Score (ES). The KS statistic seeks a single maximum cumulative deviation and is therefore most sensitive when the pathway genes accumulate toward one extreme of the ranked gene list. However, actual biological regulation is often more complex than the idealized concordant scenario. While some studies argue that pathway-associated genes tend to exhibit concordant expression changes and recommend separate enrichment analysis for up- and downregulated genes to avoid signal cancellation [13], this approach may overlook biologically meaningful bidirectional regulation patterns [14, 15]. Moreover, in many cases, the pathway activity changes may involve only a small subset of genes rather than sweeping transcriptional changes across the entire pathway [4, 16]. In such scenarios, the KS statistic may fail to accumulate sufficient signal and detect biologically meaningful enrichment patterns.

To enhance detection power and robustness across diverse scenarios, we introduce an improved GSEA method, **in**tegral-based GSEA (inGSEA). inGSEA introduces a novel enrichment score based on the Anderson-Darling (AD) [17] weighted integral statistic. The AD statistic integrates squared deviations over the entire distribution and assigns heavier weights to the tails. The integral over the distribution effectively accumulates bidirectional signals, and the tail-weighting improves sensitivity to enrichment patterns driven by a small subset of highly ranked genes. While a recent study explored the AD statistic for variance-based GSA by rank-ing samples [18], inGSEA incorporates it into the gene-ranking paradigm of GSEA. inGSEA employs the Cauchy combination [19, 20] to aggregate KS and AD statistics. The combination provides robustness to various enrichment patterns by balancing the complementary strengths of both statistics. When the concordant assumption holds, the KS statistic remains highly sensitive; otherwise, the AD statistic is more sensitive. To overcome the resolution limits imposed by finite permutations, inGSEA utilizes the generalized gamma distribution to approximate the empirical null distribution, substantially improving computational efficiency and *p*-value resolution. Taking advantage of these methodological advancements, we implement the inGSEA framework as a user-friendly web-based software tool, empowering researchers to perform rapid interactive pathway analysis within their web browsers (https://amss-stat.github.io/inGSEA).

## 2 Method

### 2.1 Standard GSEA Framework

Consider an expression dataset with *N* total genes and a predefined gene set *S* containing *N*_*H*_ genes. All *N* genes are ranked according to a correlation metric *r*_*j*_ (such as signal to noise ratio) that reflects the association between gene expression and the phenotype. This yields an ordered gene list *L* = (*g*_1_, *g*_2_, …, *g*_*N*_). GSEA evaluates whether the members of set *S* are randomly distributed throughout *L* or significantly enriched at either extreme. Such enrichment reflects that the predefined gene set, as a whole, exhibits differential expression between the two biological phenotypes. For each rank *i* ∈ {1, …, *N*}, the weighted empirical cumulative distribution of genes inside and outside the set *S* are defined as:

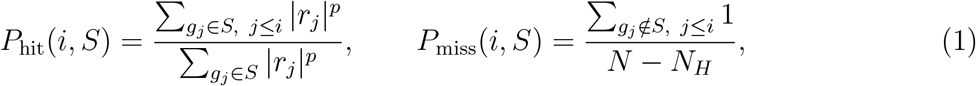

where *p* is a weighting parameter and is typically set to 1 [4].

The standard enrichment score is defined as

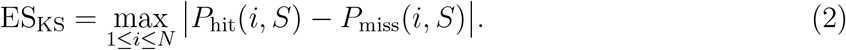

This enrichment score is essentially a Kolmogorov-Smirnov type statistic. To estimate statistical significance, the phenotype labels are randomly permuted to generate an empirical null distribution. The *p*-value *p*_KS_ is calculated as the proportion of scores obtained from phenotype permutations at least as extreme as the observed ES_KS_.

### 2.2 Enrichment Score Based on the Weighted Integral Statistic

The standard enrichment score based on *L*_*∞*_ norm is effective for strongly concordant signals where genes are largely enriched at one end of the ranked list. It may overlook sparse or bidirectional enrichment because it evaluates only the largest local deviation, ignoring the global discrepancy. The discrepancy between *P*_hit_ and *P*_miss_ is often distributed across multiple ranks without producing a single prominent peak, then the *L*_*∞*_ norm may fail to reach statistical significance even if the total evidence of pathway enrichment is substantial. To capture the distributed differences, the *L*_2_ norm is preferable.

We propose a weighted integral enrichment score ES_AD_ based on the Anderson-Darling (AD) test [17]. The continuous formulation of the AD statistic *A*^2^ integrates the weighted squared discrepancy with respect to the pooled distribution Φ(*t*):

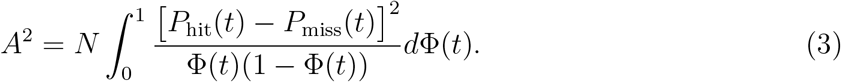

Evaluate this integral over the discrete ranks *i* ∈ {1, …, *N −* 1} and define AD enrichment score as:

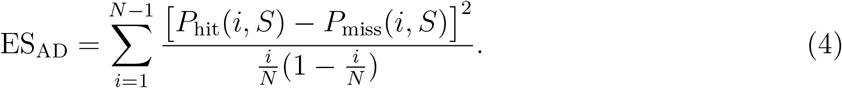

The *p*-value *p*_AD_ is calculated as the proportion of scores obtained from phenotype permutations at least as extreme as the observed ES_AD_.

ES_AD_ provides a biologically intuitive solution for detecting complex enrichment patterns. The integral statistic captures the cumulative evidence of pathway perturbation across the entire gene list, rather than relying on a single isolated peak. The numerator of ES_AD_ retains the correlation weighting |*r*_*j*_| in *P*_hit_ of GSEA, while the denominator normalizes the variance of empirical distribution. It ensures that genes with larger phenotype associations contribute more to the running sum, and discrepancies are evaluated comparably across all ranks, magnifying critical tail signals while penalizing central noise.

For sparse signals, the variance weight effectively magnifies hits at the extremes that would otherwise fail to exceed the background noise in the KS test. For bidirectional signals, the summation aggregates discrepancies from both upregulated and downregulated gene clusters into a unified measure, whereas the classic maximum score would discard all complementary evidence outside the single largest peak.

### 2.3 Cauchy Combination Method and Multiple Testing Corrections

To leverage the complementary strengths of ES_KS_ (which remains highly powerful under the concordant assumption) and ES_AD_ (which excels at detecting sparse or bidirectional signals), we combine their respective *p*-values using the Cauchy combination test [19, 20]. Let *p*_KS_ and *p*_AD_ denote the *p*-values obtained from the permutation tests. The Cauchy combined statistic is constructed as:

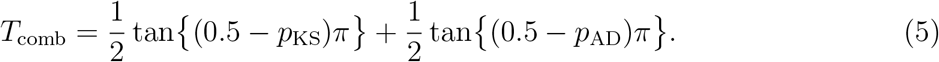

Under the null hypothesis of no enrichment, the combined *p*-value is calculated as:

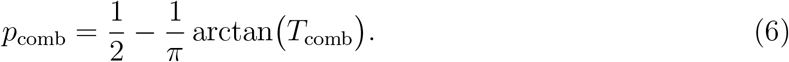

The choice of the Cauchy combination is motivated by two critical factors. First, it provides a robust solution to the dependency structure between the underlying tests. Because both statistics are evaluated on the identical ranked gene list and the same gene set, *p*_KS_ and *p*_AD_ are often correlated. Traditional methods like Fisher’s method require either strictly independent *p*-values or explicit modeling of the dependency structure. The Cauchy combination is theoretically proven to control the false positive rate (FPR) under arbitrary dependency structures [19, 20].

Second, the Cauchy combination possesses a sensitivity to the minimum *p*-value among the combined tests. In the extreme tails of the distribution, *T*_comb_ behaves similarly to a minimum function rather than an average [19, 20]. This property is ideal for inGSEA, because it ensures that the combined result remains significant if either ES_KS_ identifies a strongly concordant signal or ES_AD_ detects a sparse or bidirectional enrichment pattern. This approach allows inGSEA to maintain high sensitivity across diverse biological scenarios whenever either test yields a sufficiently significant *p*-value.

In studies involving hundreds or more pathways, multiple testing correction is typically performed to correct for occurrence of false positives. Following the standard practice in GSEA, we adjust *p*-values based on the normalized enricnment score (NES) [4, 21, 22], which yields a False Discovery Rate (FDR) for each gene set. Consistent with the original GSEA study, gene sets with FDR < 0.25 are considered significantly enriched [4].

For each gene set *S*, the normalized AD enrichment score NES_AD_ is calculated as:

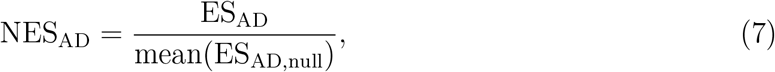

where ES_AD,null_ represents the AD enrichment scores obtained from the null distribution via phenotype permutations, and mean(*·*) reprents calculating the average. The normalization places AD enrichment scores from gene sets of different sizes on a common scale and enables the FDR estimation procedure in GSEA [4]. Since ES_AD_ is always positive, it does not need to be processed separately for positive and negative values as in GSEA.

### 2.4 Empirical null distribution Approximation

A well-known bottleneck of the standard GSEA is its reliance on extensive permutations to establish an accurate null distribution. To obtain high resolution *p*-values, empirical permutation requires at least *O*(1*/p*) resamplings, making the computation of extreme tail probabilities very expensive. Analytical approximation of the null distribution has been proposed as an effective acceleration strategy [23]. Specifically, blitzGSEA demonstrated that the null distribution of the standard enrichment score ES_KS_ can be approximated by a gamma distribution [11].

Since AD enrichment score ES_AD_ is formulated as a weighted *L*_2_ norm involving squared deviations, its null distribution exhibits distinct tail characteristics compared to the *L*_*∞*_-based enrichment score ES_KS_. The gamma approximation for GSEA is no longer optimal. We conducted a comprehensive numerical study to identify the most suitable parametric family for ES_AD_ by evaluating several candidates, including the lognormal, chi-square, gamma and generalized gamma distributions [24], across KEGG legacy pathway database. The generalized gamma distribution is a highly flexible family that subsumes several common distributions [24, 25], such as the gamma, Weibull, and exponential distributions, as special cases. Based on the Bayesian Information Criterion (BIC) [26], the generalized gamma distribution provided the best fit among all candidate distributions (see Table 1). Furthermore, simulations on real-world datasets confirmed that the generalized gamma approximation maintains well-calibrated false discoveries (see Section 3.1 and Figure 1).

**Table 1:** Comparison of BIC values for fitting the null distributions of ES_AD_. Values represent the mean across all tested 137 KEGG legacy pathways containing at least 20 genes, where the BIC for each gene set was first averaged over 1,000 independent repetitions (each employing 1,000 permutations). Lower values indicate superior fit.

| Distribution | BIC |
| --- | --- |
| **Generalized Gamma** | **16653.6217** |
| Log-normal | 16654.5962 |
| Gamma | 16673.7549 |
| Weibull | 16759.3960 |

**Figure 1:**
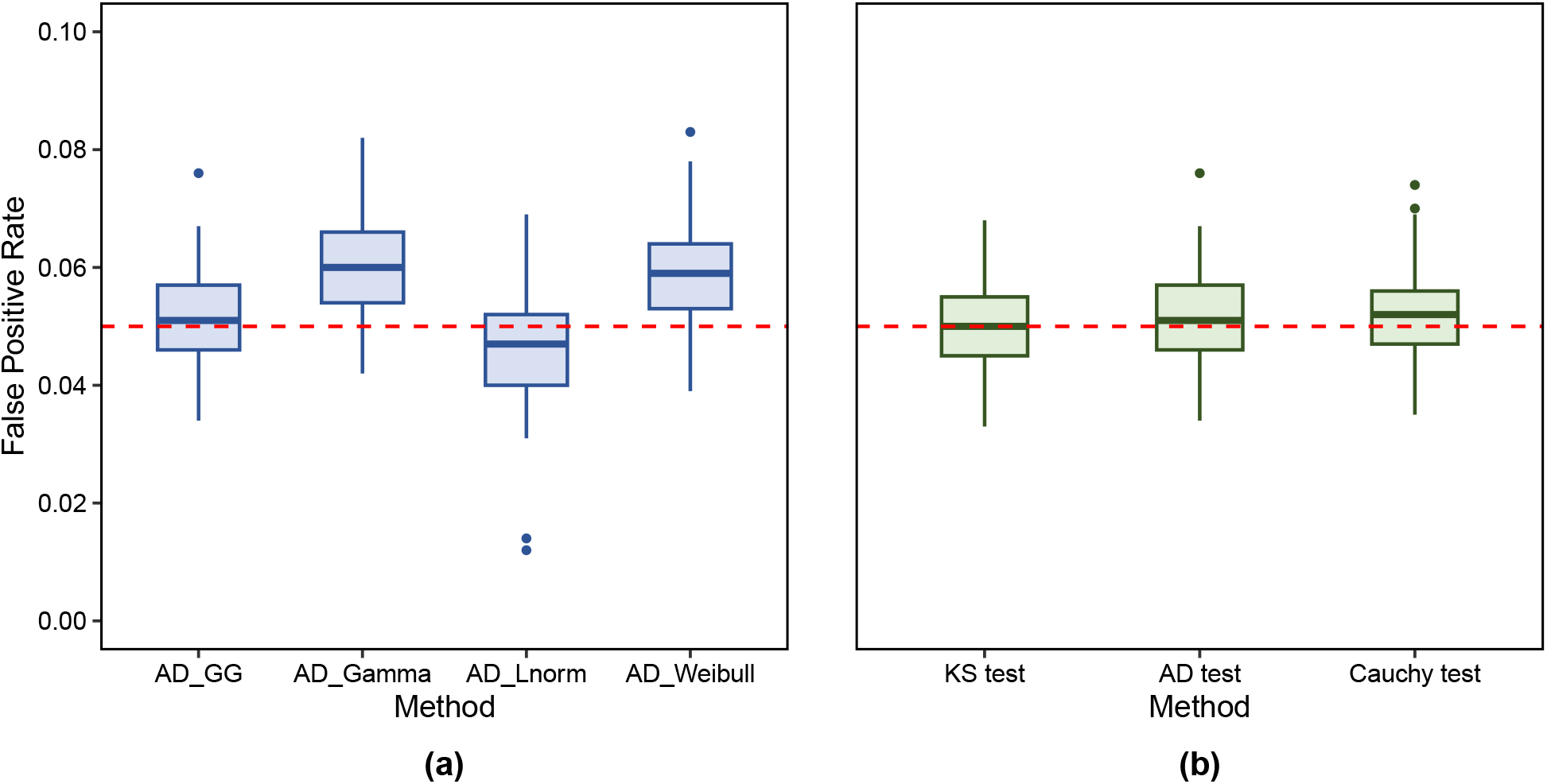
Evaluation of false positive rates across KEGG legacy pathways. (a) FPR boxplots when approximating the null distribution of ES_AD_ using different parametric distributions. (b) Comparison of FPR controlled by different methods. The horizontal dashed line indicates the nominal level of 0.05. Each boxplot represents the distribution of FPR values calculated across the 137 KEGG legacy pathways containing at least 20 genes.

### 2.5 Interactive Web-based Software

To facilitate the practical application of inGSEA, we developed a user-friendly interactive web-based software tool (https://amss-stat.github.io/inGSEA). The software is designed with a streamlined workflow: users simply drag and drop local expression datasets and gene set files (e.g., in.gmt format) into the designated input area, configure essential parameters, and initiate the analysis with a single click. The software provides comprehensive outputs, including *p*-values, FDR values, enrichment scores, and interactive enrichment plots and charts.

The tool leverages modern client-side web technologies to provide a high-performance analytical tool without the need for local software installation or programming expertise. The key features of the inGSEA software are summarized below:

#### Privacy via Client-Side Computation

Unlike traditional web tools that require uploading sensitive genomic data to a remote server, inGSEA performs all statistical computations locally within the user’s web browser. No expression data is transmitted to servers, ensuring complete confidentiality and compliance with data privacy regulations.

#### Efficiency and Real-Time Feedback

Benefiting from the generalized gamma approximation described in Section 2.4, inGSEA significantly reduces the computational burden. Analysis of hundreds of pathways typically completes within an hour on a standard personal computer. The interface features a real-time progress bar and estimated time-to-completion, keeping the user informed throughout the computational process.

#### Dynamic Interactivity and Visualization

The inGSEA tool offers an advanced interactive interface for exploring results. Analysis outputs are presented in a dynamic table where pathway names are hyperlinked, allowing users to jump directly to source databases, such as MSigDB [5, 6], for detailed biological context. Furthermore, the enrichment plots are fully interactive. By hovering the curosr over or clicking specific segments of the enrichment plot, users can identify individual genes and their corresponding rank metrics. This real-time exploration helps researchers bridge the gap between statistical significance and biological interpretation. The output also includes NES charts for all pathways to clearly show the main enrichment directions and strengths.

#### Compatibility and Open Accessibility

The tool inGSEA is fully compatible with the standard GSEA workflow and MSigDB. To comply with copyright restrictions of MSigDB, users are encouraged to download MSigDB files (.gmt) themselves or use custom gene sets. inGSEA is accessible via a public URL (https://amss-stat.github.io/inGSEA), and its source code is hosted on GitHub (https://github.com/amss-stat/inGSEA). Detailed user guides and step-by-step tutorials are provided in the software’s homepage and GitHub README.

## 3 Results

### 3.1 False positive rate

We conducted a comprehensive simulation study to evaluate the FPR. The data were generated from a real-world transcriptomic dataset “Gene regulation and DNA damage in the ageing human brain” (AHB dataset) [27] and the KEGG legacy pathway database. The simulation was based on a multivariate normal distribution, whose mean vector and covariance matrix were estimated from the AHB dataset. We employed shrinkage estimation to compute a robust covariance matrix. Under the null hypothesis, simulated samples were randomly assigned to case (age *>* 42) and control (age *≤* 42) [27] groups.

The procedure was repeated for 1,000 independent experiments, with each analysis involving 1,000 permutations. We calculated the FPR at a significance level of *α* = 0.05 for the standard GSEA (KS test), AD test and Cauchy combination methods. Specifically for ES_AD_, we calculated its *p*-value using both the permutation method and several parametric approximations, including the gamma, lognormal, Weibull, and generalized gamma (GG) distributions.

We generated boxplots of the FPR calculated for each KEGG legacy pathway containing at least 20 genes. As depicted in Figure 1(a), among the parametric approximations for the null distribution of ES_AD_, the generalized gamma distribution provided the most accurate control of the FPR. Figure 1(b) illustrates that inGSEA, including the AD test and the Cauchy combination test, effectively controlled the FPR around the nominal level of 0.05.

### 3.2 Statistical power

We conducted a comprehensive power analysis to assess our methods in detecting pathway enrichment under various scenarios. The simulation was based on the AHB dataset and the Hallmark pathway database, leveraging the multivariate normal distribution.

Recognizing that pathway-specific attributes such as size, gene-gene correlations, and biological function can influence detection power, we focused on three representative pathways with distinct sizes and high gene overlap with the expression data: Hallmark Hedgehog Signaling (small, with 33 genes), Hallmark Peroxisome (medium, with 84 genes), and Hallmark Myc Targets V1 (big, with 181 genes).

For each pathway, we systematically injected artificial signals into the case group by modifying the mean expression levels of a subset of pathway genes. Three signal injection paradigms were simulated to mimic different biological phenomena:

1. Concordant signal: A coordinated upregulation was simulated by randomly selecting 80% of the pathway genes and increasing their mean expression in the case group. The magnitude of the increase was set to 0.3, 0.4, and 0.5 times each gene’s population standard deviation. This aligns with the basic assumptions and motivations of the original GSEA literature.
2. Sparse signal: To model scenarios where only a small core subset of genes is activated, we simulated a sparse signal by randomly selecting only 20% of the pathway genes and upregulating them by the same effect sizes (0.3, 0.4, and 0.5 standard deviations).
3. Bidirectional signal: Reflecting complex biological processes where some genes are activated while others are repressed, we simulated a bidirectional signal. We randomly selected 40% of the pathway genes for upregulation and a distinct 40% for downregulation, with both subsets’ expression levels shifted by 0.3, 0.4, and 0.5 standard deviations from the mean.

For each combination of pathway, signal type, and effect size, we performed 1,000 independent simulation experiments. In each experiment, GSEA was conducted with 2,000 permutations. We focused on the methods validated in the FPR analysis: *p*-values are calculated using the generalized gamma approximation. Empirical power was defined as the proportion of the 1,000 simulations in which the signal-injected pathway was detected as significant at an *α* level of 0.05.

Figure 2 summarizes the results of the power simulations. In the concordant signal paradigm, all evaluated methods demonstrated high statistical power, with the AD test and Cauchy combination methods performing comparably to or slightly better than the standard GSEA. The superior sensitivity of our methods became more pronounced in the more challenging signal patterns. Specifically, the AD test achieved an approximately 2%-12% increase in power for detecting sparse signals and a more substantial 9%-40% gain in the bidirectional signal scenario compared to the standard GSEA. These findings align with the analysis in the Methods section, confirming that inGSEA is more powerful and robust at detecting complex enrichment patterns.

**Figure 2:**
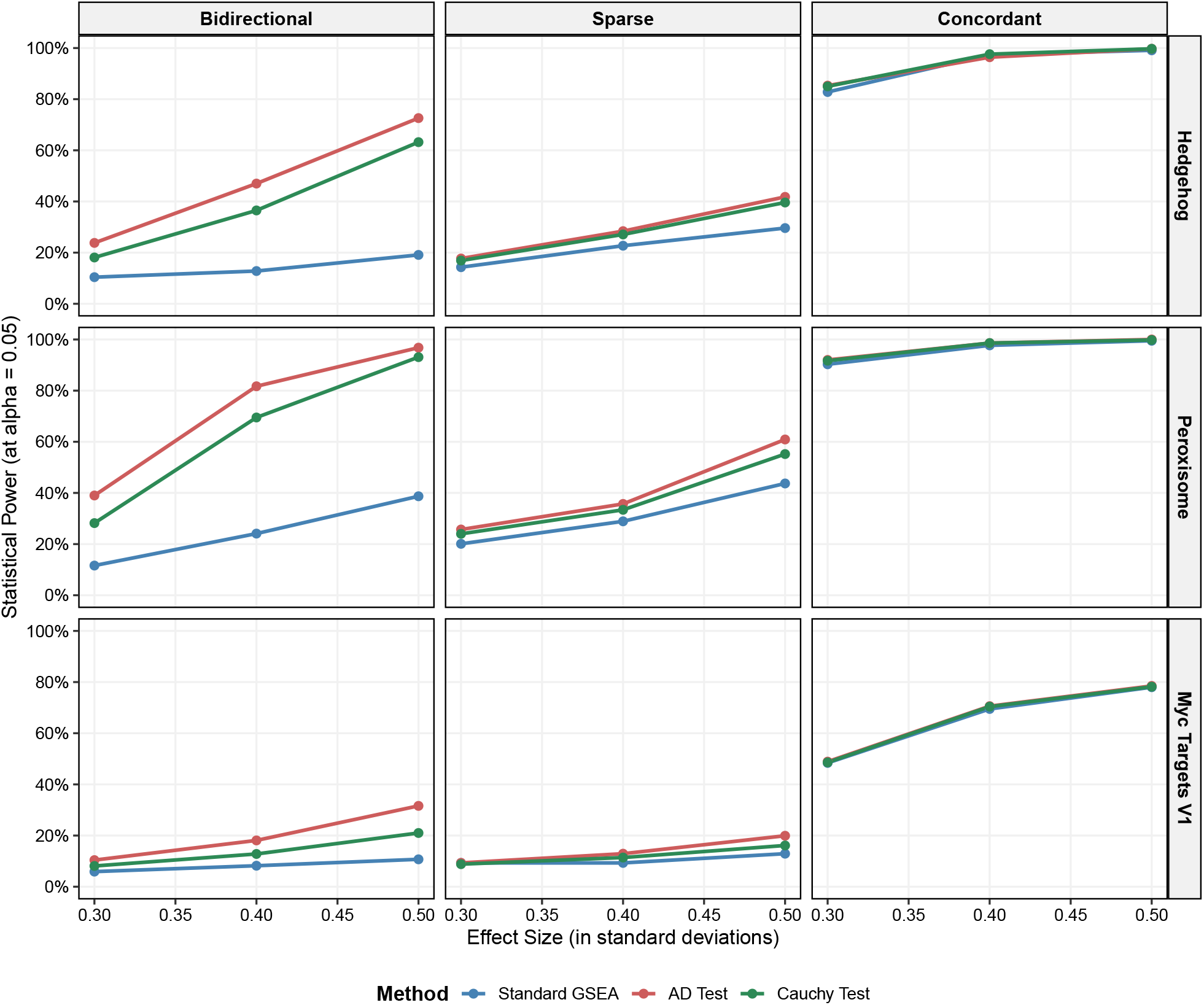
Statistical power comparison of the standard GSEA and inGSEA across different signal paradigms and pathway sizes. The 3 × 3 grid displays three signal paradigms (columns: bidirectional, sparse, and concordant) across three representative pathways of varying sizes (rows: Hedgehog Signaling with 33 genes, Peroxisome with 84 genes, and Myc Targets V1 with 181 genes). The x-axis represents the signal strength (effect sizes of 0.3, 0.4, and 0.5 standard deviations), and the y-axis represents the empirical power evaluated at a nominal level of *α* = 0.05. Each data point is calculated based on 1,000 independent simulation runs with 2,000 permutations. Different colored curves represent the standard GSEA and inGSEA methods (AD test and Cauchy combination test). The null distribution of AD test is approximated using the generalized gamma distribution.

### 3.3 Application to real data

We applied our method to four transcriptomic datasets representing diverse biological contexts: the AHB dataset [27], GSE157103 (SARS-CoV-2 infection) [28], GSE12452 (nasopharyngeal carcinoma) [29, 30], and GSE32323 (colorectal cancer) [31]. We utilized multiple authoritative pathway databases, including Hallmark, KEGG legacy and Reactome. To evaluate and contrast the discovery power of inGSEA against the standard GSEA, we employed a nominal significance level of *α* = 0.05 and FDR threshold of < 0.25, as recommended by the original GSEA.

Across the tested datasets and databases, inGSEA demonstrated higher discovery power. Our method identified 144 novel pathways that showed evidence of pathway enrichment but were missed by the standard GSEA. Conversely, 58 pathways identified by the standard GSEA were not detected by inGSEA. 243 pathways were identified by both methods. We plotted corresponding overlap Venn diagrams (see Figure 3). Figure 4 shows the distribution of significant pathways across different databases. Notably, in GSE157103 dataset (SARS-CoV-2 infection), the set of significant pathways identified by the standard GSEA is almost entirely a subset of those identified by inGSEA across all three databases. It indicates that inGSEA retains the discovery capacity of the standard GSEA while capturing additional enrichment signals. In the vast majority of cases, inGSEA identified more significant pathways than GSEA.

**Figure 3:**
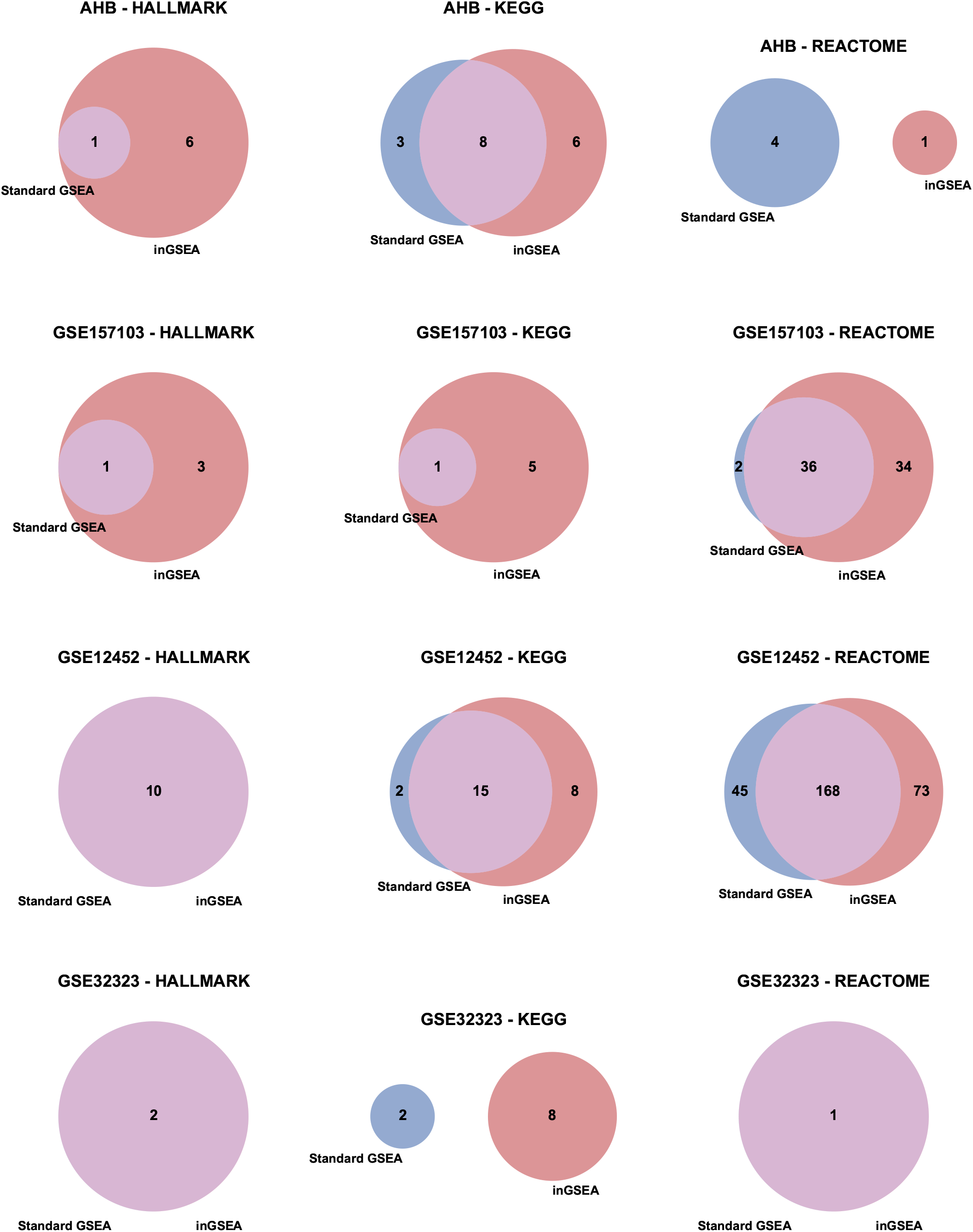
Venn diagrams displaying the overlap of significant pathways (*p*-value < 0.05 and FDR < 0.25) identified by the standard GSEA and inGSEA. The figure comprises 12 subfigures corresponding to each dataset-database combination. Red circles represent inGSEA, while blue circles represent the standard GSEA. The numbers in the non-overlapping and overlapping regions indicate the count of unique and shared significant pathways, respectively.

**Figure 4:**
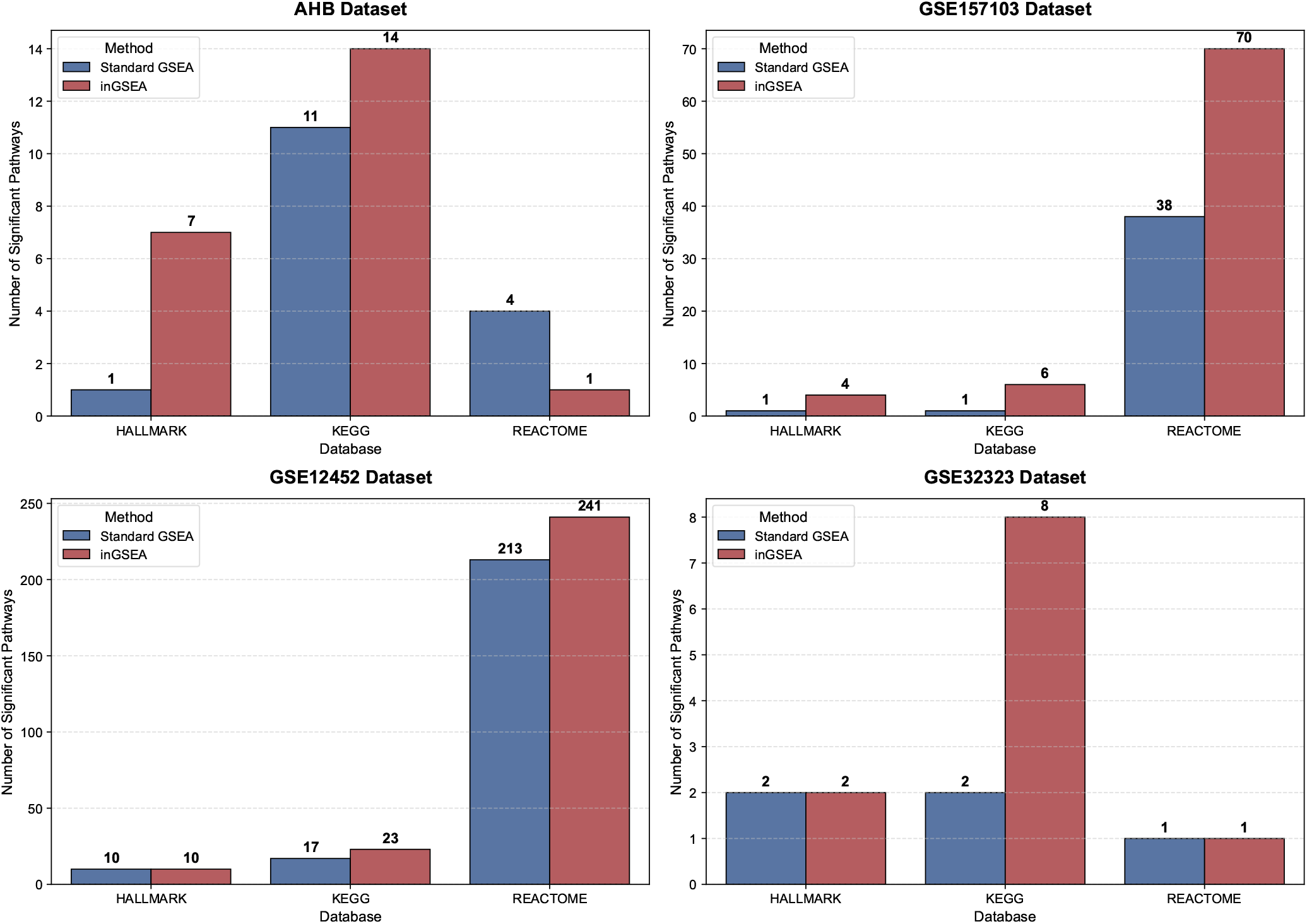
Comparison of the number of significant pathways (*p*-value < 0.05 and FDR < 0.25) identified by standard GSEA and inGSEA. The figure consists of four subfigures, each representing one of the four tested expression datasets. Within each subfigure, the x-axis displays the three pathway databases (Hallmark, KEGG, and Reactome). Blue and red bars denote the standard GSEA and inGSEA, respectively, with the exact counts of identified pathways annotated on top of each bar.

Detailed results for benchmark database Hallmark are summarized in Table 2, providing the *p*-values, FDR values and documented biological evidence for the novel pathways. Most of these new findings are supported by independent biological studies, suggesting that they represent genuine pathophysiological signals rather than statistical artifacts.

**Table 2:**
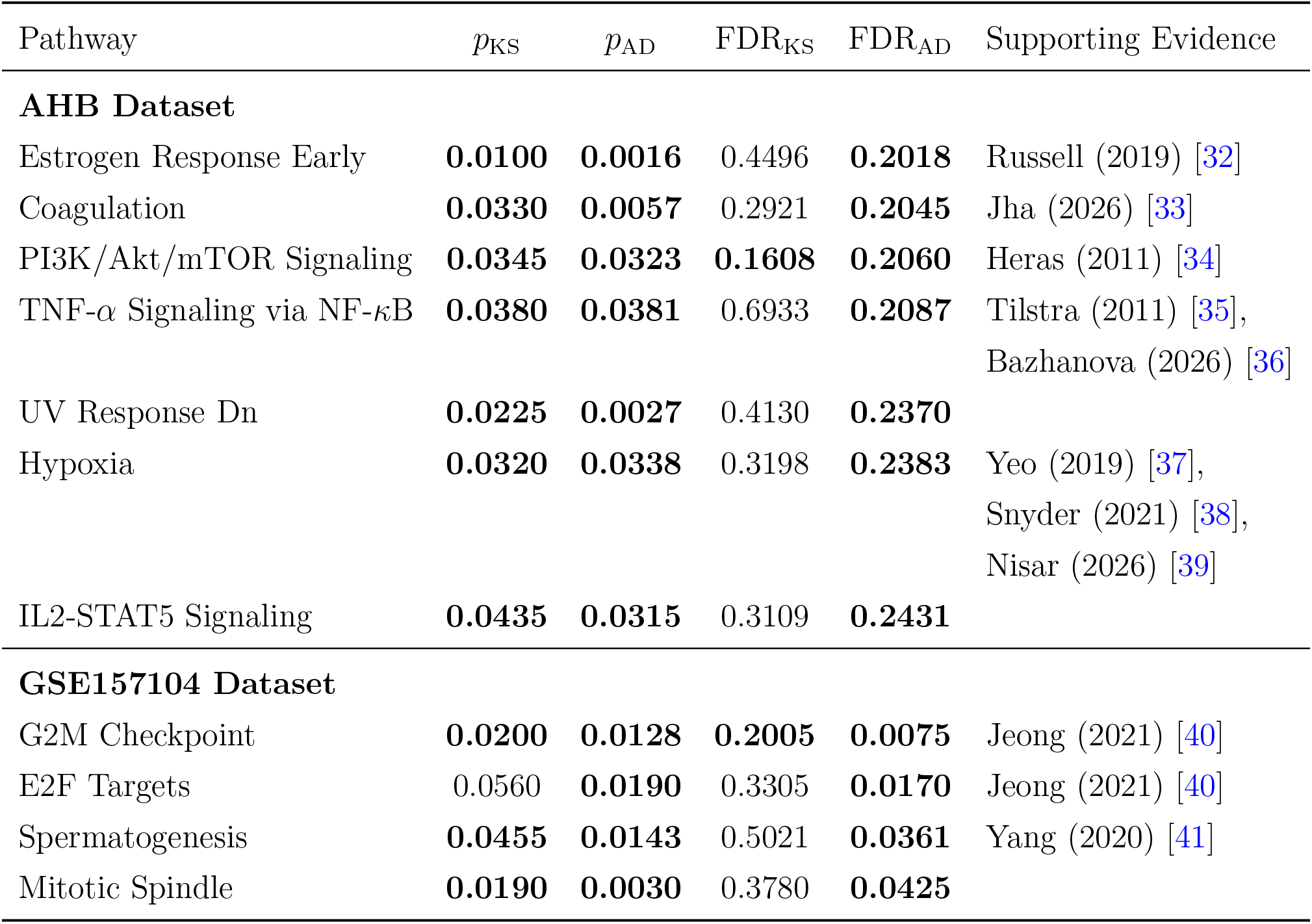
Representative significant pathways from Hallmark database discoveries across AHB and GSE157104 datasets. Significant *p*-values (< 0.05) and FDR values (< 0.25) are highlighted in **bold**. Pathway names are simplified for clarity. *p*_KS_ and FDR_KS_ correspond to the standard GSEA, while *p*_AD_ and FDR_AD_ correspond to inGSEA.

In the AHB dataset, which profiles transcriptional changes in the aging human brain, the Estrogen Response Early pathway provides a representative example. Estrogen signaling is known to play a protective role in the brain, and its decline is associated with cognitive and physiological aging [32]. Under the standard GSEA, this pathway yielded a nominal *p*-value of 0.01, but failed to survive the multiple testing correction with an FDR_KS_ of 0.4496, potentially overlooking a biologically relevant pathway. In contrast, inGSEA detected significant enrichment for this pathway (*p*_AD_ = 0.0016, FDR_AD_ = 0.2018).

In the GSE157104 dataset associated with SARS-CoV-2 infection, the contrast is even more pronounced for the E2F Targets pathway. This pathway regulates crucial cell cycle and proliferative programs, which are frequently dysregulated during active viral replication [40]. The standard GSEA did not identify this pathway as significant, yielding an insignificant *p*-value of 0.056 and an FDR_KS_ of 0.3305. In contrast, inGSEA detected significant enrichment for the same pathway (*p*_AD_ = 0.019, FDR_AD_ = 0.017).

The advantage of inGSEA is visually clarified by the enrichment plot (see Figure 5). A strong transcriptional signal is evident at the leading edge, with a substantial enrichment of pathway genes concentrated within the top 400 ranked genes. Following the initial enrichment, the running enrichment score fluctuates within a stable range between 0.52 and 0.58 across a wide span of the ranked list, eventually reaching its maximum deviation at rank 3,275. Notably, the peak is broad and relatively flat rather than sharp. Because the KS statistic in the standard GSEA relies heavily on the absolute maximum deviation (the peak), it lacks the sensitivity to resolve such a late, broad peak, especially when preceded by prolonged fluctuations. In contrast, inGSEA, utilizing the weighted integral statistic, integrates deviations over the entire ranked list. It successfully captures both the prominent early enrichment before rank 400 and the sustained moderate signal, resulting in a statistically significant detection.

**Figure 5:**
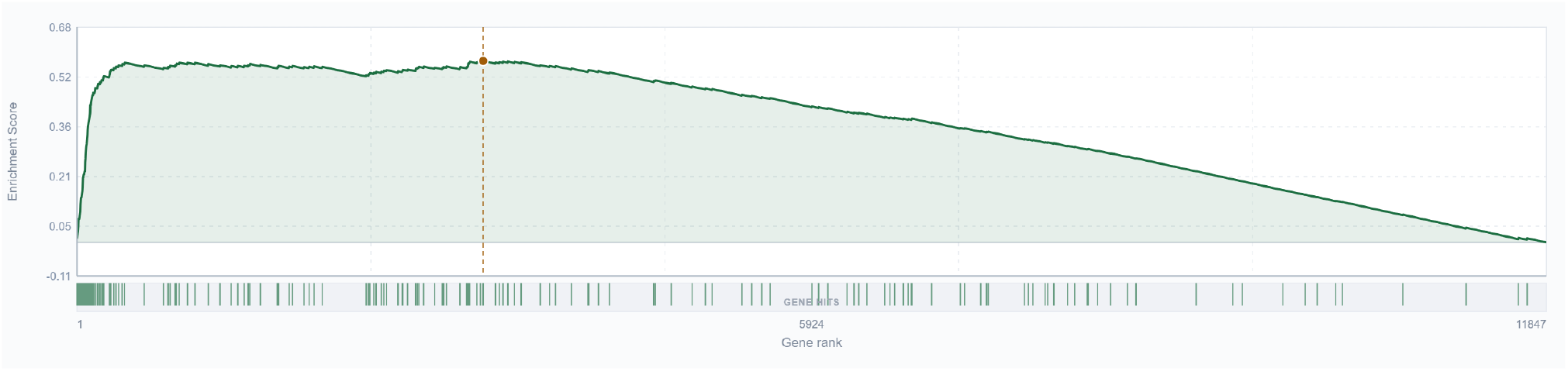
Enrichment plot for the E2F Targets pathway in the GSE157104 dataset. The top panel shows the running enrichment score, and the bottom panel (vertical green bars) indicates the positions of the pathway genes along the ranked gene list. The red dashed line indicates the location of the gene corresponding to the peak enrichment score.

## 4 Discussion

We present inGSEA, an improved method designed to overcome the concordant assumption constraints of the standard GSEA and the computational bottlenecks of permutation testing. By introducing the weighted integral enrichment score and coupling it with the standard GSEA via a Cauchy combination, inGSEA provides a powerful and robust gene set enrichment analysis method. It preserves the power of the standard GSEA for concordant pathways while improving detection power for complex signals. By shifting the paradigm from exhaustive permutation to the generalized gamma approximation, inGSEA effectively reduces computing time.

The primary innovation of inGSEA lies in adopting the weighted integral statistic instead of the KS statistic to calculate the enrichment score ES_AD_. Unlike the KS statistic, which focuses only on the maximum single deviation, the AD statistic integrates the squared discrepancy over the entire distribution. It allows the AD statistic to more effectively capture dispersed signals across the whole gene list. Furthermore, the AD statistic assigns higher weights to genes at both tails of the ranked list, making it more sensitive to crucial genes. As demonstrated by simulation results, inGSEA exhibits higher power when detecting bidirectional or sparse enrichment patterns. Real data analysis also confirms that less ideal, non-concordant pathways indeed exist in specific datasets. For instance, in the E2F pathway, while the main peak of the enrichment plot is not sharp, pathway genes are clearly enriched at both extremes. These findings strongly support inGSEA as a robust and powerful method for gene set analysis.

We developed a user-friendly web-based software tool. Rather than requiring local installation or programming skills, the inGSEA tool features a streamlined drag-and-drop interface. The tool utilizes client-side computation, meaning all analyses are performed directly within the user’s web browser. The software provides dynamic interactivity: users can explore hyperlinked results, visualize NES charts to determine main enrichment directions, and interact with enrichment plots to identify specific genes and their rank metrics. It bridges the gap between statistical significance and biological interpretation while remaining fully compatible with standard GSEA workflows and MSigDB.

Despite these strengths, our study has several limitations that warrant consideration. First, the AD enrichment score accounts for genes at both tails of the ranked list. While this improves statistical power, it introduces potential interpretability challenges. In practice, we recommend investigating the specific enrichment direction of significant pathways with NES charts and enrichment plots. Our interactive software greatly facilitates this process, allowing users to quickly identify enriched genes at both tails. One potential scenario is that pathway genes at both extremes are not co-regulated but display distinct positive and negative peaks on the enrichment curve. Although such exploratory results are not statistical false positives, their biological interpretation requires caution and is highly dependent on the biological context and the quality of the gene sets.

Second, the current distribution approximation method still requires a baseline number of permutations. When analyzing a massive number of pathways, stringent multiple testing correction necessitates highly precise *p*-values, which implies that thousands of permutations or more are still required. Although the distribution approximation strategy drastically reduces the computational burden compared to pure permutation testing, it still demands considerable computing time under extreme multiple testing scenarios. Future research could explore hybrid strategies that combine permutation acceleration algorithms with distribution approximation to further enhance computational efficiency.

There are limitations regarding the scope of application. Currently, the method is not directly applicable to single-sample enrichment analysis, nor is it optimally designed for highly sparse data. Future work will focus on extending inGSEA to broader contexts, such as adapting the integral statistic for single-sample profiling or resolving sparsity issues for single-cell data. In conclusion, inGSEA provides a powerful and robust improvement over the standard GSEA, effectively reducing the reliance on the concordant assumption and enhancing the detection power. We anticipate that inGSEA will become a widely adopted tool in the field of gene set analysis, helping researchers uncover novel enriched pathways and deepening our understanding of complex pathway functions.

## Data availability

All data used in this study are publicly available. Gene expression datasets were obtained from the NCBI Gene Expression Omnibus (GEO) under accession numbers GSE157103, GSE12452, and GSE32323. Additional datasets were collected from previously published studies as cited in the main text. Pathway definitions were obtained from the MSigDB database (version 2026.1).

## Code availability

inGSEA is available at https://github.com/amss-stat/inGSEA.

## Acknowledgements

This research was supported by National Natural Science Foundation of China (Grant No. 12325110, 12288201).

